# Improved vectors for retron-mediated CRISPR-Cas9 genome editing in *Saccharomyces cerevisiae*

**DOI:** 10.1101/2024.08.06.606807

**Authors:** Tara N. Stuecker, Stephanie E. Hood, Julio Molina Pineda, Sonali Lenaduwe, Joshua Winter, Meru J. Sadhu, Jeffrey A. Lewis

## Abstract

*In vivo* site-directed mutagenesis is a powerful genetic tool for testing the effects of specific alleles in their normal genomic context. While the budding yeast *Saccharomyces cerevisiae* possesses classical tools for site-directed mutagenesis, more efficient recent CRISPR-based approaches use Cas ‘cutting’ combined with homologous recombination of a ‘repair’ template that introduces the desired edit. However, current approaches are limited for fully prototrophic yeast strains, and rely on relatively low efficiency cloning of short gRNAs. We were thus motivated to simplify the process by combining the gRNA and its cognate repair template in *cis* on a single oligonucleotide. Moreover, we wished to take advantage of a new approach that uses an *E. coli* retron (EcRT) to amplify repair templates as multi-copy single-stranded (ms)DNA *in vivo*, which are more efficient templates for homologous recombination. To this end, we have created a set of plasmids that express Cas9-EcRT, allowing for co-transformation with the gRNA-repair template plasmid in a single step. Our suite of plasmids contains different antibiotic (Nat, Hyg, Kan) or auxotrophic (*HIS3, URA3*) selectable markers, allowing for editing of fully prototrophic wild yeast strains. In addition to classic galactose induction, we generated a β-estradiol-inducible version of each plasmid to facilitate editing in yeast strains that grow poorly on galactose. The plasmid-based system results in >95% editing efficiencies for point mutations and >50% efficiencies for markerless deletions, in a minimum number of steps and time. We provide a detailed step-by-step guide for how to use this system.

## Introduction

*In vivo* site-directed mutagenesis is a powerful genetic tool for testing the effects of specific alleles in their normal genomic context. The budding yeast *Saccharomyces cerevisiae* is a powerful genetic model due to strong ‘classical’ tools for genome modification based on high rates of homologous recombination of donor DNA. There have been a number of methods for generating targeted genome edits in yeast (reviewed in Fraczek et al., 2018), with markerless editing of essential genes being the historic challenge.

The *delitto perfetto* method (Storici & Resnick, 2006) was among the first breakthroughs in producing *in vivo* site-directed mutations in yeast. The method relies on insertion of a selectable marker into the gene of interest, followed by transformation of a 140 bp double-stranded oligonucleotide that contains homology to the insertion site with the targeted mutation near its center. Homologous recombination results in swapping out the marker cassette with the allele of interest, with counter selection being used to identify transformants that have lost the original marker. This method results in mutagenesis efficiencies of ∼20% (Storici & Resnick, 2006) for non-essential genes. Essential genes can be mutated too by inserting the marker cassette directly downstream of the gene of interest, but with lower mutagenesis efficiencies.

More recently, the advent of CRISPR-based methods for genome editing has revolutionized the construction of yeast strains. There are a number of different approaches to CRISPR-based editing, with the most common using a nuclease (usually *Streptococcus pyogenes* Cas9) that is targeted by a guide RNA (gRNA) to a specific region of the genome.

The genomic region of interest must contain a protospacer-adjacent motif (PAM, NGG sequence) that immediately follows the DNA sequence targeted by the gRNA (see Tian et al., 2017 for a review on the CRISPR machinery). The original CRISPR-based tools for site-directed mutagenesis in yeast used gRNA and Cas9 plasmids with auxotrophic markers for selection that were co-transformed with ∼90-nt double-stranded oligonucleotides containing the repair template (DiCarlo et al., 2013; Hu et al., 2018; Laughery et al., 2015). The efficiency of point mutations ranged from >50% (Hu et al., 2018) to nearly 100% (DiCarlo et al., 2013; Laughery et al., 2015). However, a downside for those methods is that they require generation of auxotrophic strains prior to transformation with CRISPR-editing plasmids. More recently developed systems include Cas9 and gRNA plasmids with antibiotic resistance markers (Mans et al., 2015; Vyas et al., 2018), allowing editing of fully prototrophic wild strains. Likewise, the “insert, then replace” strategy of *delitto perfetto* has been combined with CRISPR to increase efficiencies (Lutz et al., 2019), though this still requires an added step of marker integration near the editing site plus the use of ds oligonucleotides or PCR products as repair templates. Thus, we were motivated to simplify the process by using a strategy originally designed for CRISPR libraries where each gRNA and its cognate repair template are paired in *cis* on a single oligonucleotide (Sadhu et al., 2018). An additional benefit of this approach is that cloning short gRNAs has been a challenge (Antony et al., 2022), so using a longer oligonucleotide that combines the gRNA and repair template should yield higher cloning efficiencies.

An exciting recent advance in the design of yeast CRISPR libraries is the Cas9 Retron precISe Parallel Editing via homologY (CRISPEY) method (Sharon et al., 2018). The CRISPEY method uses an *E. coli* “retron” to amplify repair templates as multi-copy single-stranded (ms)DNA *in vivo*. Retrons are natural DNA elements that consist of a reverse transcriptase gene (EcRT) and a template that the EcRT acts upon to generate msDNA (Inouye & Inouye, 1992). Generating a chimeric fusion of the retron, guide RNA, and repair template leads to amplification of the repair template as msDNA when the reverse transcriptase is expressed, which in a test case increased editing efficiency for a point mutation to nearly 100% (Sharon et al., 2018).

While the CRISPEY system was originally designed for generating thousands of edits in parallel, the ease of gRNA and repair template design, cloning, and editing efficiency would make it a valuable tool for targeted editing. However, the originally designed CRISPEY system relies on integration of the Cas9 and EcRT into the genome at a *his3Δ1* location, which requires one or two extra strain construction steps depending on whether the strain of interest already has the *his3Δ1* allele. To enable editing without having to first integrate Cas9-EcRT into the genome, we created a set of centromeric (CEN) plasmids that express Cas9-EcRT, allowing for co-transformation with the gRNA/repair template plasmid in a single step. Additionally, our suite of plasmids contains different antibiotic (Nat, Hyg, Kan) or auxotrophic (*HIS3, URA3*) selectable markers, allowing for editing of fully prototrophic wild yeast strains. Finally, we generated a β-estradiol inducible version of the CRISPEY system to facilitate editing in yeast strains that grow poorly on galactose. The plasmid-based system results in >95% editing efficiencies for point mutations and >50% efficiencies for markerless deletions, in a minimum number of steps and time. We provide a detailed step-by-step guide for how to use this system.

## Materials and methods

### Strains, media and reagents

Strains used in this study are listed in Table 1, and all media components and reagents are listed in Table S1. *E. coli* was grown in lysogeny broth (LB: 1% tryptone, 0.5% yeast extract, 0.5% NaCl) containing 100 μg/ml ampicillin at 37°C with 270 rpm orbital shaking. Yeast growth media included YPD (1% yeast extract, 2% peptone, 2% dextrose), YP-Gal (1% yeast extract, 2% peptone, 2% galactose), and synthetic complete (Sherman, 2002) media lacking either uracil (SC-Ura) or histidine (SC-His). When antibiotics were used with SC media, 0.1% monosodium glutamate was used as a nitrogen source. Antibiotics were used at the following final concentrations: nourseothricin at 100 μg/ml, G418 at 200 μg/ml, hygromycin B at 300 μg/ml. Yeast were propagated in liquid media at 30°C with 270 rpm orbital shaking. When added to the media, β-estradiol was at a concentration of 1 µM. Yeast were transformed using the method of (Gietz & Schiestl, 2007). For auxotrophic selection, cells were resuspended in 2% glucose and plated directly on the appropriate SC dropout plates. For antibiotic selection, cells were resuspended in YPD and plated onto non-selective media (either YPD or SC complete as appropriate) and incubated 16-24 hours before replica printing to antibiotic plates. Haploid derivatives were generated by deletion of *HO*, followed sporulation and dissection, and then PCR screening (Illuxley et al., 1990) to identify MATa strains.

**Table 1.**
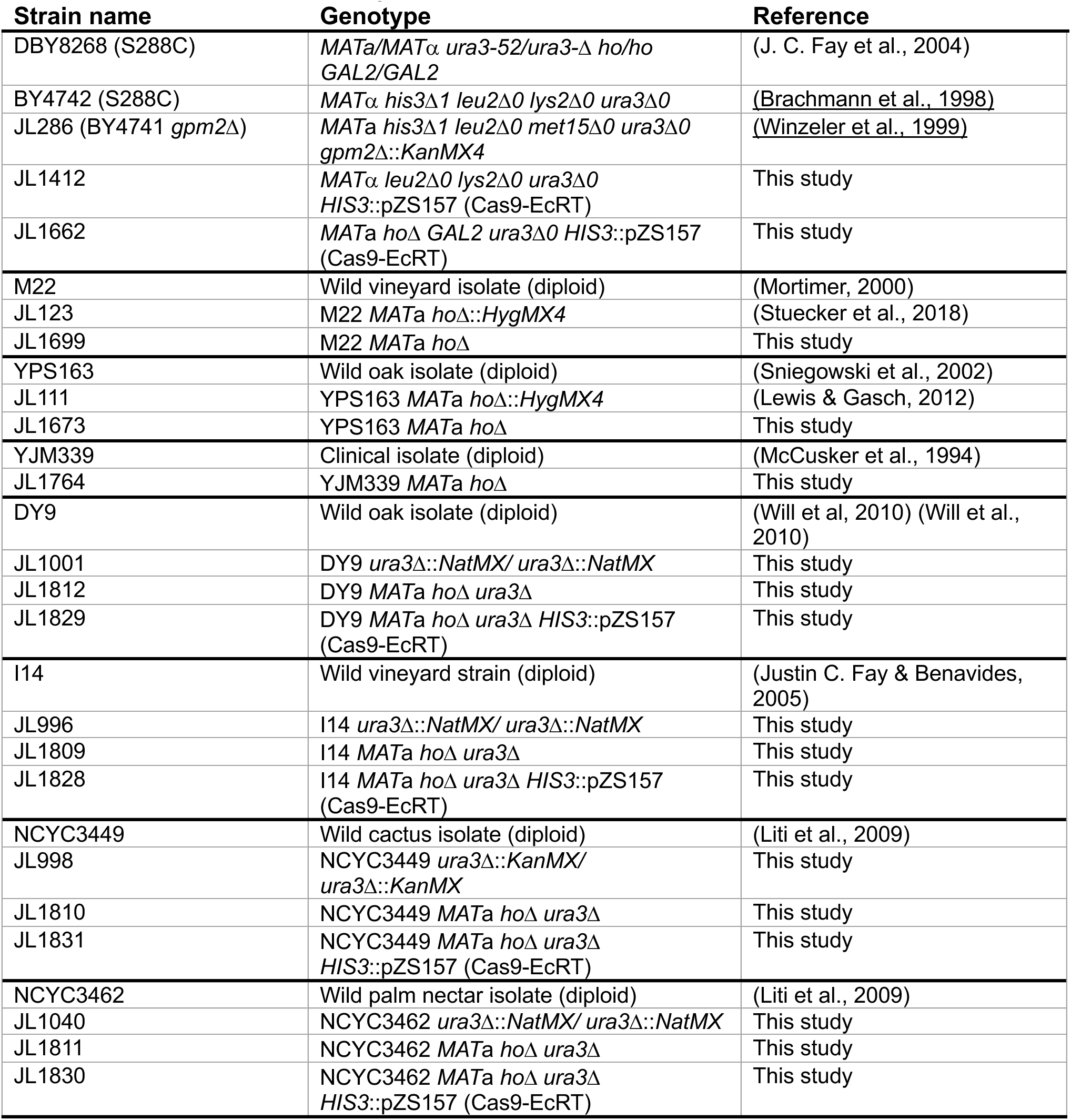
Strains used in this study.

### General plasmid construction methods

All primers used in this study were purchased from Integrated DNA Technologies (IDT) and are listed in Table S2. PCR amplification conditions listed in Table S3. Restriction enzyme digests were incubated at 37°C overnight (16 – 20 h). DNA fragments were ligated at a 5:1 molar ratio of insert to vector using 5 units of T4 ligase, 5% PEG4000, and a 20 h incubation with 30 second cycling between 10°C and 30°C. PCR products and linearized vectors were either column-purified with the Zymo DNA Clean & Concentrator-5 Kit or gel-purified with the Zymoclean Gel DNA Recovery Kit, as indicated, and eluted with nuclease-free water. Cloning products were transformed into either chemically-competent *E.coli* DH5α or Stellar^TM^ competent cells (Takara) via heat shock using the manufacturer’s instructions for Stellar cells. Plasmids were isolated from *E. coli* cultures using the Promega Wizard® Plus Miniprep DNA purification kit. To isolate plasmids from yeast cells, 5-10 ml of yeast culture was pelleted and resuspended in 250 μl of the Wizard® Plus kit Cell Resuspension Buffer. Cells were vortexed for 5 min with 100 ul of glass beads at room temp, then beads were allowed to settle for 10 min. Supernatant was transferred to a sterile microcentrifuge tube and plasmids were purified using the Wizard® Plus kit starting at step 2. Cloned constructs were sequenced via long-read Oxford Nanopore sequencing (Plasmidsaurus).

### Cas9-EcRT plasmid construction

All plasmids expressing Cas9 and the Ec86 reverse transcriptase (hereafter abbreviated as Cas9-EcRT) are listed in Table 2, and full plasmid sequences can be found in File S1. pCas9-EcRT-GAL-Hyg was constructed by amplifying and inserting the Ec86 reverse transcriptase (EcRT) and *S. pyogenes* Cas9 (Cas9) from the pZS157 CRISPEY RT/Cas9 integration plasmid (Sharon et al., 2018) into the CEN plasmid pAG26 (Goldstein & McCusker, 1999), which carries *URA3* and HygMX markers. EcRT and Cas9 are arranged in pZS157 CRISPEY RT/Cas9 opposite of the bidirectional *GAL1-GAL10* promoter, and were thus amplified as a single PCR product (p*GAL1*::Cas9::t*CYC1* p*GAL10*::EcRT::t*GAL10*) using Cas9-EcRT-pAG26_PfoI_F and Cas9-EcRT-pAG26_Mph1103I_R primers. The resulting PCR product was digested with *Dpn*I to remove the pZS157 template plasmid. pAG26 plasmid was linearized by simultaneously digesting with *Pfo*I and *Mph*1103I. Both the Cas9-RT PCR product insert and linearized pAG26 vector were column-purified and then fused together using the Takara In-Fusion® HD Cloning Kit to create pCas9-EcRT-GAL-Hyg.

**Table 2.**
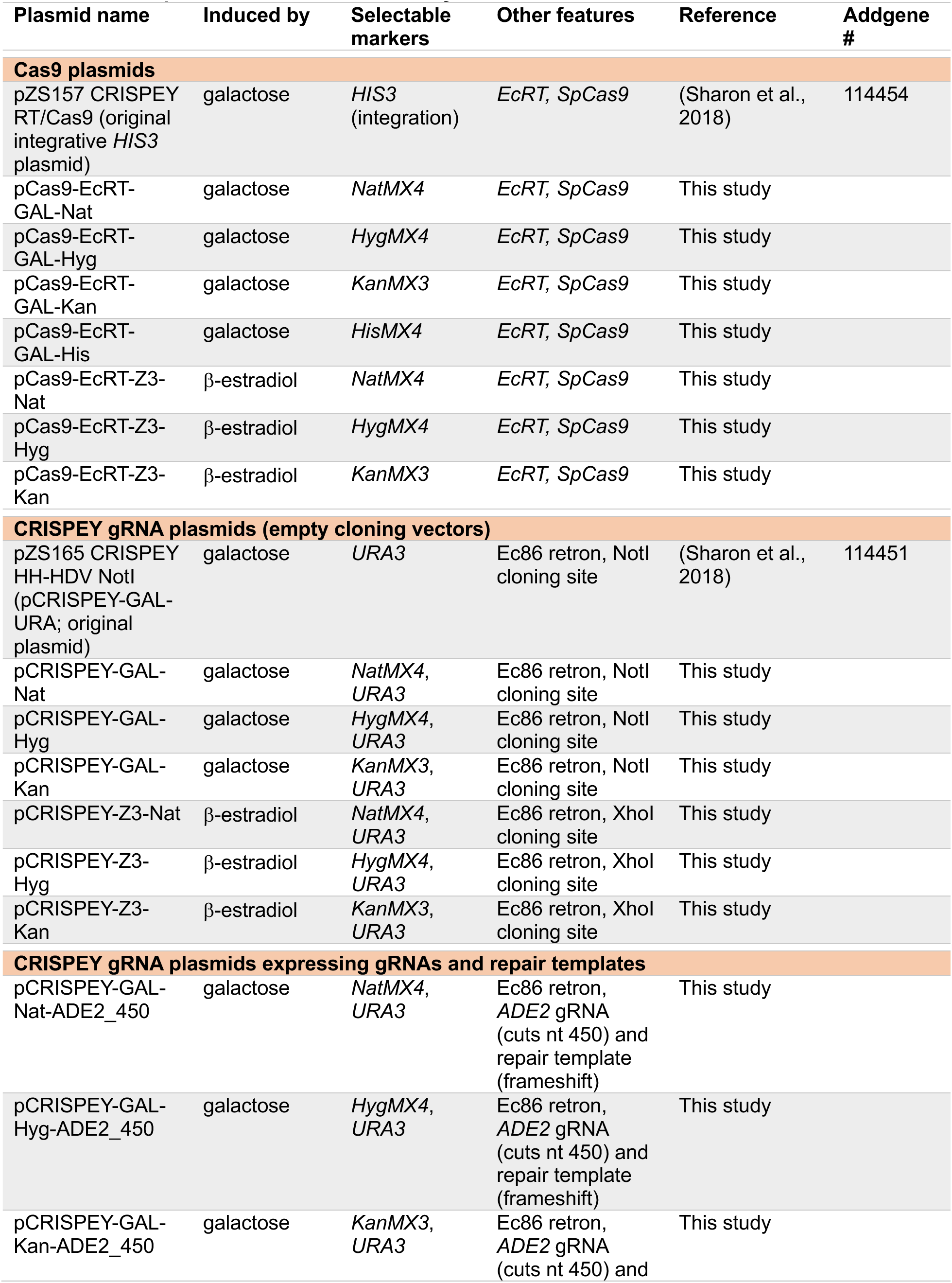
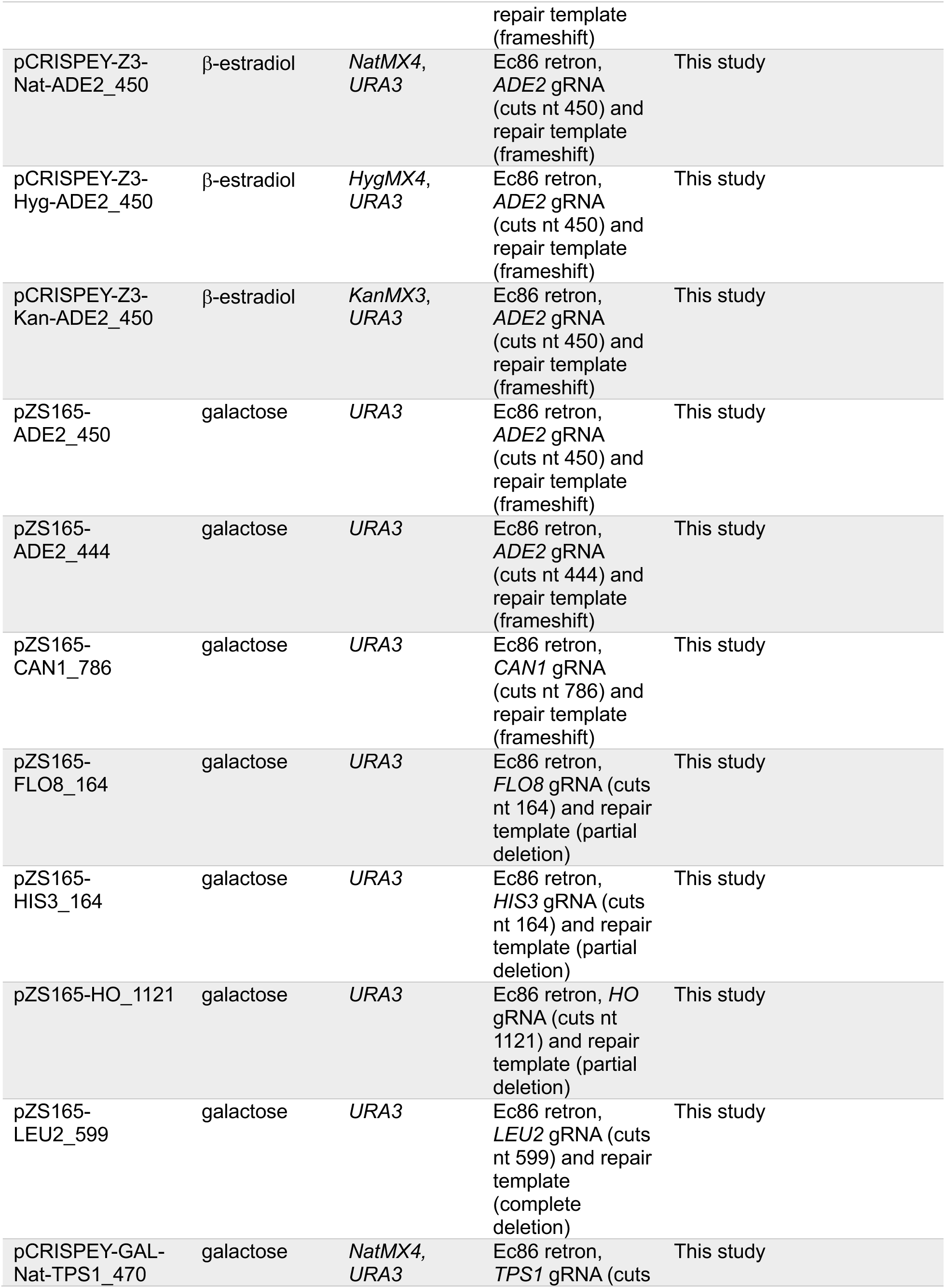
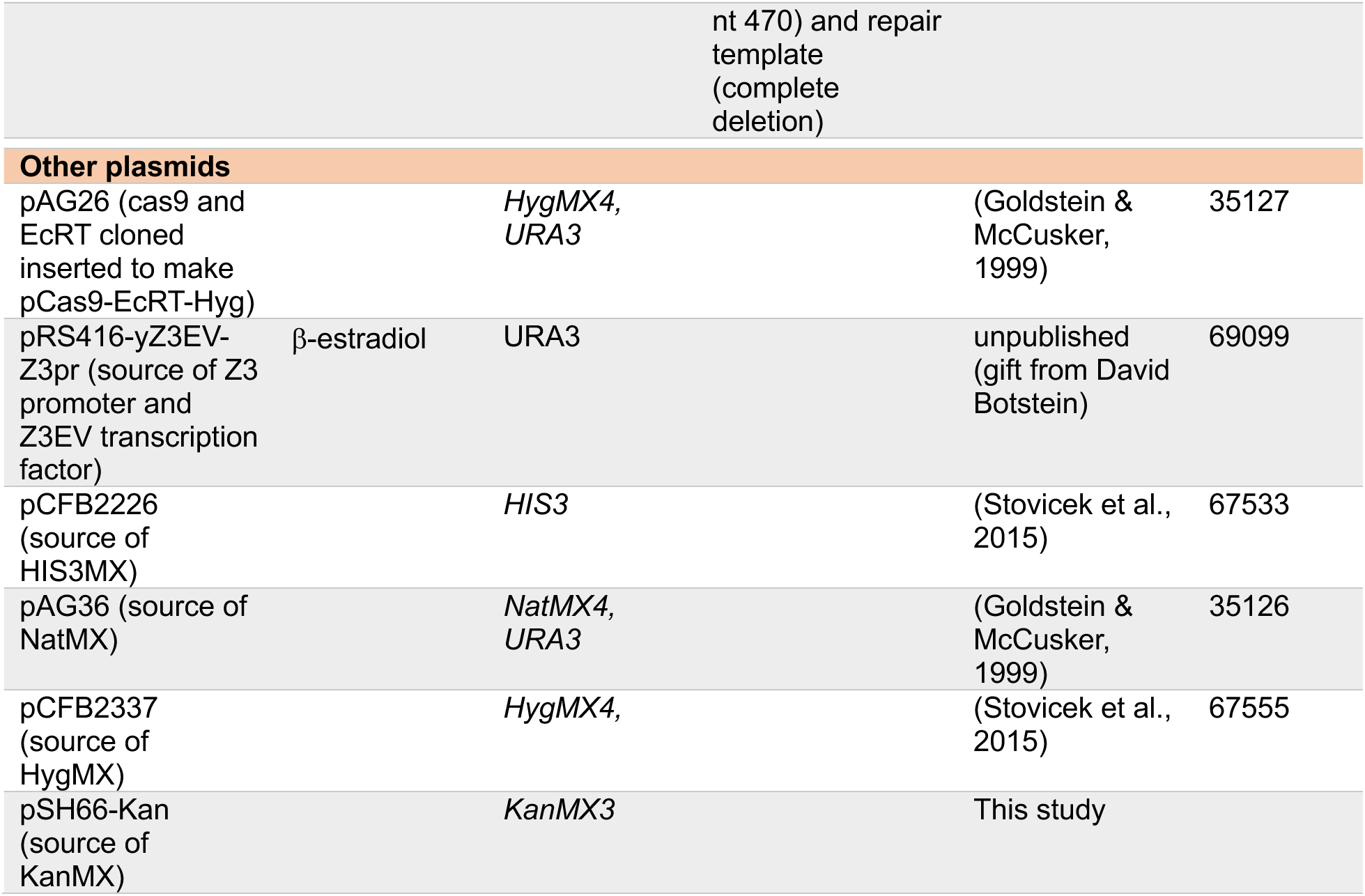
List of plasmids used in this study.

pCas9-EcRT-GAL-Nat and pCas9-EcRT-GAL-Kan plasmids were created by using yeast homologous recombination to replace the HygMX cassette with PCR products containing either a KanMX or NatMX (amplified from JL286 or JL111, respectively). The KanMX or NatMX cassettes were amplified using PR78 and PR79 primers that anneal to *TEF* promoter and terminator sequences common to all MX cassettes (Goldstein & McCusker, 1999). pCas9-RT-GAL-Hyg was transformed into BY4742, followed by transformation of the KanMX or NatMX PCR products. Plasmids from transformants that lost hygromycin B resistance (HygMX-) and gained either G418 resistance (KanMX+) or nourseothricin resistance (NatMX+) were isolated from yeast transformed into *E. coli* DH5α, and then re-isolated and sequenced.

The pCas9-EcRT-GAL-His plasmid was created by replacing the HygMX cassette in pCas9-EcRT-GAL-Hyg with a His3MX cassette. The pCFB2226 plasmid was digested with *Eco*91I and *Mss*I and the 1.25-kb fragment containing His3MX was gel-purified. The pCas9-EcRT-GAL-Hyg vector was digested with *Eco*91I and *Oli*I to excise the HygMX cassette. The 10.0 kb vector fragment (lacking HygMX) was gel-purified and ligated to the His3MX fragment with T4 ligase. The resulting product was transformed into *E. coli* and isolates were screened by diagnostic restriction digest. Plasmids with the correct digest patterns were verified by transformation into yeast cells, followed by phenotypic screens (His+ and Hyg-) and whole plasmid sequencing.

Estradiol-inducible Cas9-EcRT plasmids were each created in two steps: replacement of the bidirectional *GAL1-GAL10* promoter with the Z_3_ promoter, followed by insertion of the Z_3_EV transcription factor into a non-coding region of the vector. The bidirectional *GAL1-GAL10* promoter for each of the pCAS9-EcRT-GAL plasmids was replaced with the Z_3_ promoter by Gibson cloning (Gibson et al., 2009). The Z_3_ promoter was amplified from pRS416-yZ3EV-Z3pr (a gift from David Botstein) using CloneAmp^TM^ HiFi PCR Premix according to manufacturer’s instructions with the following conditions: initial denaturization at 98°C for 1 min followed by 35 cycles of 98°C for 10 sec, 55°C for 15 sec, 72°C for 1 min with a final extension step at 72°C for 5 min. Linear, promoter-less Cas9-EcRT vectors were created by PCR with Z3pr_CAS_VF and Z3pr-CAS_VR primers using Herculase II Fusion DNA polymerase according to manufacturer instructions for amplifying large products. The amplification conditions were an initial denaturization at 95°C for 2 min followed by 10 cycles of 95°C for 15 sec, 55°C for 20 sec, 68°C for 5.5 min, then another 20 cycles where the 68°C extension step time is increased incrementally by 20 sec per cycle followed by a single extension step at 72°C for 8 min after the 30 cycles. Both vector and Z_3_ promoter insert PCR products were digested with *Dpn*I to remove template plasmids, column-purified, then fused together using the NEBuilder^®^ HiFi DNA Assembly cloning kit according to manufacturer instructions. Next, the Z_3_EV transcription factor was cloned into the ‘intermediate’ plasmids containing the Z_3_ promoter driving Cas9 and EcRT. The Z_3_EV transcription factor was amplified from pRS416-yZ3EV-Z3pr using Z3TF_CAS_Ins_F and Z3TF_CAS_Ins_R primers with CloneAmp^TM^ HiFi PCR Premix. The PCR conditions were identical to those used to amplify the Z_3_ promoter fragment with the exception of the extension step, which was increased to 2 min to accommodate the larger product size (∼2 kb). The ‘intermediate’ plasmids were linearized with *Mph*1103I, and the PCR fragment containing the Z_3_EV transcription factor was cloned into the *Mph*1103I site using the NEBuilder^®^ HiFi DNA Assembly cloning kit.

### CRISPEY gRNA plasmid construction

All CRISPEY plasmids for expressing gRNAs and homologous repair templates are listed in Table 2. The pCRISPEY-GAL plasmids were constructed by inserting an MX antibiotic resistance cassette from a donor plasmid into the original pZS165 CRISPEY HH-HDV vector (Sharon et al., 2018). The NatMX cassette was excised from pAG36 (Goldstein & McCusker, 1999) via *Bam*HI-*Eco*RI double digestion. *Bgl*II-*Sac*I double digests were used to excise the HygMX cassette from pCFB2337 (Stovicek et al., 2015) and the KanMX cassette from pSU66. The DNA fragments containing each antibiotic resistance cassette were gel purified, and the 3’ recessed (aka ‘sticky’) ends were filled in by incubation with 3 units of T4 DNA polymerase and 0.2 mM dNTPs in NEB Buffer 2.1 for 5 min at 37°C. The resulting blunt-ended fragments were column purified and ligated into 50 ng *Pvu*II-linearized pZS165 CRISPEY HH-HDV vector (at a 5:1 molar ratio of insert to vector) using T4 ligase. Following transformation of the ligation reaction into *E. coli*, plasmids from individual clones were isolated and screened for inserts via diagnostic restriction digest. The presence of each selectable antibiotic marker was confirmed by transforming final plasmids into yeast and verifying antibiotic resistances, followed by whole plasmid sequencing. The gRNAs and repair templates were cloned into CRISPEY plasmids as described in the user protocol in the Results and Discussion section.

To create the pCRISPEY-Z3 plasmids, the unique *Not*I cloning site in the pCRISPEY-GAL vectors had to be replaced with a unique *Xho*I restriction sequence due to the presence of a *Not*I site in the Z3 promoter. Each pCRISPEY-GAL plasmid was linearized by *Not*I digest and column purified. The *Xho*I site was inserted into each linearized vector by Gibson cloning with the XhoI_swap_CRISPEY oligo using the NEBuilder® HiFi DNA Assembly cloning kit according to manufacturer instructions. Intermediate plasmids containing the GAL promoter and *Xho*I cloning site were amplified by PCR with Z3_CRISP_VF and Z3_CRISP_VR primers to linearize vectors and remove the GAL promoters. The Z_3_ promoter and Z_3_EV transcription factor (including ACT1 promoter and 3’UTR) were amplified as a single fragment from pRS416-yZ3EV-Z3pr with Z3_CRISP_Ins_F and z3_CRISP_Ins_R primers. Fragments were assembled by Gibson cloning with the NEBuilder® HiFi DNA Assembly kit and transformed into *E. coli*. Plasmids were verified by diagnostic restriction digest followed by whole plasmid sequencing.

### Gene editing experiments

Editing efficiency was measured systematically using an *ADE2* reporter, where loss-of-function mutations cause the cell to accumulate purine precursors that turn colonies red when oxidized (Smirnov et al., 1967). The reporter plasmid (pCRISPEY-ADE2) contains a gRNA that directs Cas9 to cleave at nucleotide position 450 of the *ADE2* gene. This position was chosen using CRISpy-pop (Stoneman et al., 2020), which displays all possible PAM motifs for a yeast gene along with an activity ‘score’ derived using sgRNA Scorer 2.0 (Chari et al., 2017). Position 450 was chosen based on being within the first half of the *ADE2* open reading frame, and having a strong predicted gRNA activity (gRNA score of 1.34, which is above the median predicted activity. The pCRISPEY-ADE2 reporter plasmid also expresses a repair template that introduces a cytosine after position 472 of the *ADE2* gene that disrupts the PAM sequence (thus preventing further Cas9 cutting) and introduces a frameshift. Each induction experiment was done in biological triplicate on separate days to capture day-to-day variation in editing efficiencies. Each strain was transformed with one Cas9-EcRT plasmid and pCRISPEY-ADE2 plasmid. Approximately 5 unique colonies per replicate experiment were inoculated from the transformation plates into non-inducing medium (SC raffinose for galactose-inducible editing plasmids and SC glucose for estradiol-inducible editing plasmids). After 24 h growth in non-inducing media, strains were sub-cultured into induction medium (SC galactose for galactose-inducible editing plasmids and SC glucose or YPD containing 1 μM β-estradiol for estradiol-inducible editing plasmids), and were sub-cultured into fresh induction media at 24 and 48 h. For S288c, cells were diluted 30-fold for each sub-culture. Because certain wild strains grow slowly on galactose, all experiments with wild strains instead included diluting cells to a standard OD_600_ of 0.5. Editing efficiencies were measured at 0 h, 24 h, 48 h, and 72 h of induction. Serial dilutions were plated onto either YPD or SC plates, unless otherwise indicated. Plates were incubated for 3 days then the number of white and red colonies were counted, and the percent editing efficiency was calculated as the percentage of red colonies relative to the total number of colonies. For a subset of experiments, Sanger sequencing (Eurofins Genomics) was performed to calculate the percentage of *ade2* mutants with the precise frameshift mutation. For non-*ADE2* edits that were performed using this system, percent editing efficiency was calculated via phenotyping, PCR screening (for deletions), and/or Sanger sequencing (for point mutations).

## Results and Discussion

### Construction of expanded CRISPEY plasmid system

The original CRISPEY vectors were designed for parallel genome editing in a single strain background (Sharon et al., 2018). In this context, it made sense to have the genes for Cas9 and the Ec86 reverse transcriptase genomically integrated. However, this is undesirable for routine genome editing for several reasons. First, it increases the number of steps and time required to generate edits. At the very least, there is one extra step for integrating the Cas9 and EcRT, and for prototrophic strains, there is the additional step of making mutations in the *HIS3* gene to generate an auxotrophy for selection of subsequent integration. Second, genomic integration of Cas9 and the EcRT means that those genes will be induced whenever cells are growing on galactose, which likely imposes a fitness cost under that condition, and Cas9 expression even in the absence of gRNAs could lead to genomic instability (Xu et al., 2020).

Finally, because the yeast GAL induction system is the one most commonly used for over-expression studies, genomically integrated galactose-inducible Cas9 is thus non-optimal if researchers wish to analyze a genome-edited mutant’s behavior when overexpressing a gene of interest. As a solution and to simplify targeted genome editing in any strain background, we created a suite of centromeric plasmids that still allow for targeted genome editing while not relying on integration of Cas9-EcRT (Figure 1). In addition to auxotrophic markers (*URA3* or *HIS3*), the suite of plasmids also includes antibiotic resistance markers (Nat, Kan, or Hyg), thus facilitating editing in fully prototrophic wild strains of yeast.

**Figure 1.**
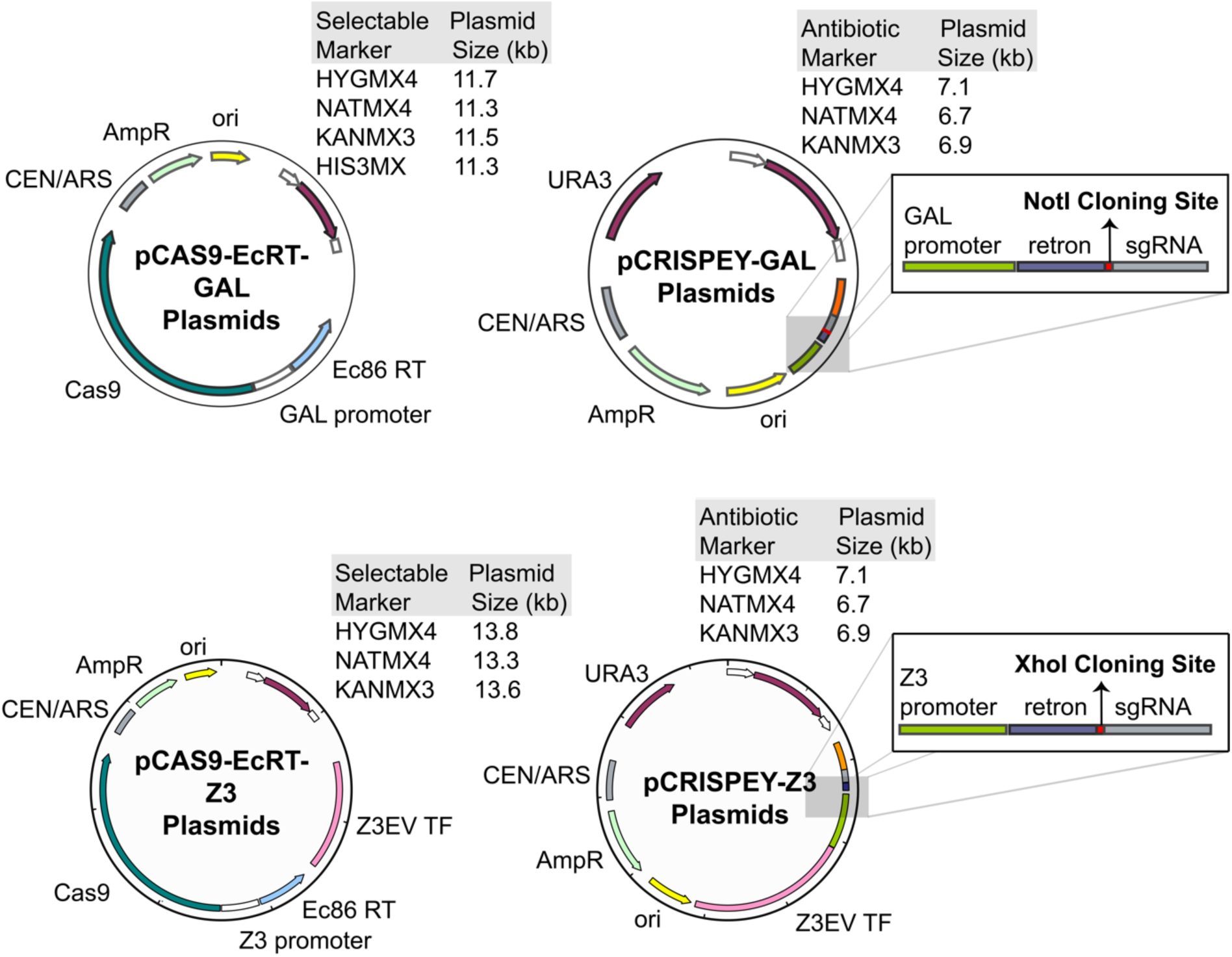
Maps of new Cas9 and CRISPEY plasmids. Two-plasmid systems for either galactose induction (A) or estradiol induction (B). The pCRISPEY plasmids contain single restriction enzyme sites for cloning (Gibson assembly), NotI for GAL (A) or XhoI for Z_3_ (B). gRNAs and their repair templates for targeted editing are synthesized as oligonucleotides and cloned into the vectors. Editing is be performed by induction from 24 – 72 hours following co-transformation of the pCRISPEY and pCAS9-Ec86 plasmids into the strain of interest.

### High genome editing efficiencies for the dual plasmid system

To test editing efficiencies, we used a gRNA and repair template that generated a frameshift mutation in the *ADE2* gene, allowing us to assess editing efficiencies for standard point mutations that would be commonly used for genomic ‘site-directed mutagenesis’ or allelic exchange experiments. *ADE2* encodes phosphoribosylaminoimidazole carboxylase (Stotz & Linder, 1990), which catalyzes the sixth step of de novo purine biosynthesis. *ade2* loss-of-function mutations accumulate P-ribosylamino imidazole (AIR), which upon oxidation turns red (Smirnov et al., 1967). Thus, the percentage of edited colonies can be determined by red-white screening. Sequencing confirmed that 100% (10/10) of the colonies contained the correct frameshift mutation at nucleotide position 472 (Table 3), consistent with Sharon and colleagues’ findings (Sharon et al., 2018) that the CRISPEY system strongly biases towards homologous recombination instead of non-homologous end joining.

**Table 3.**
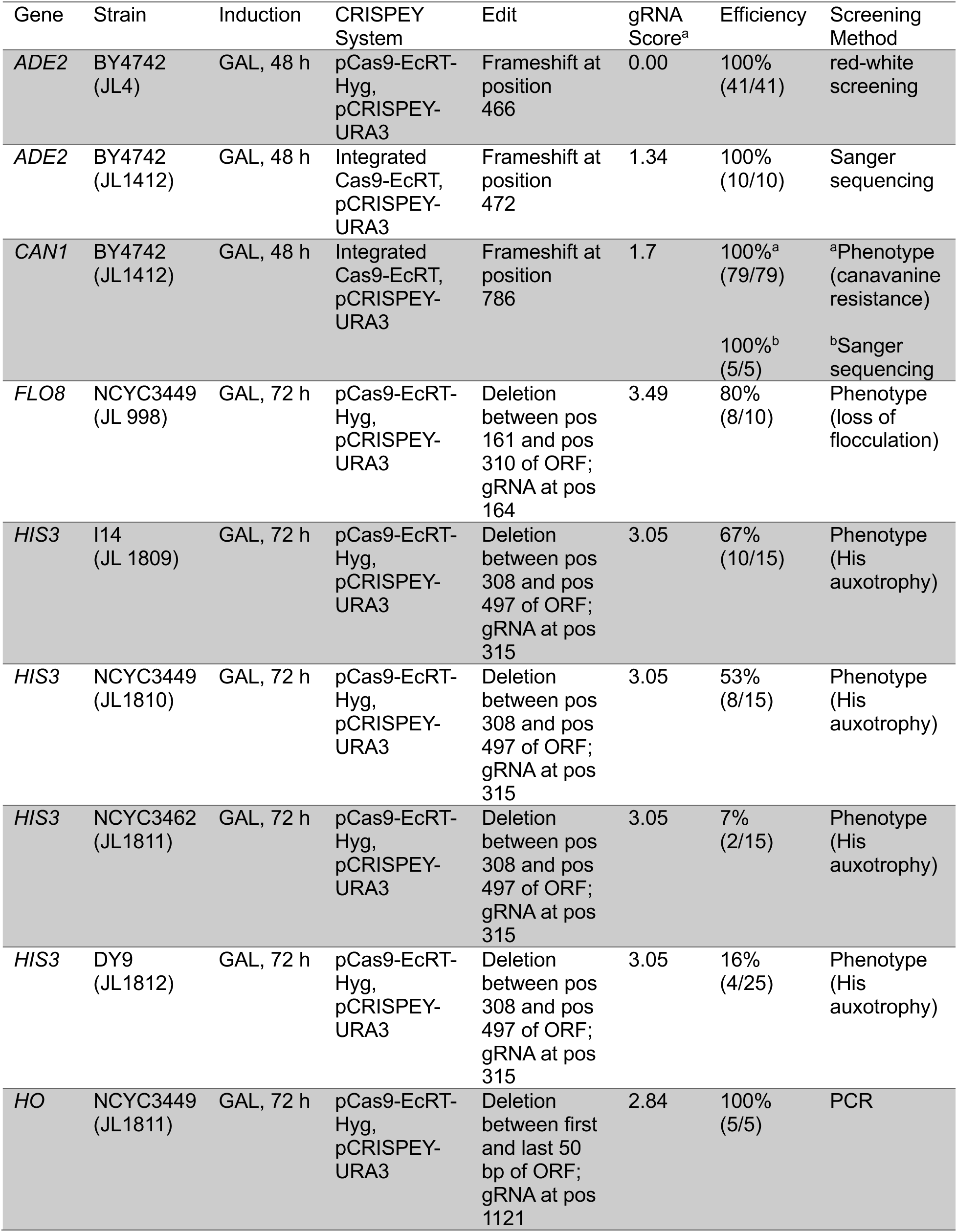
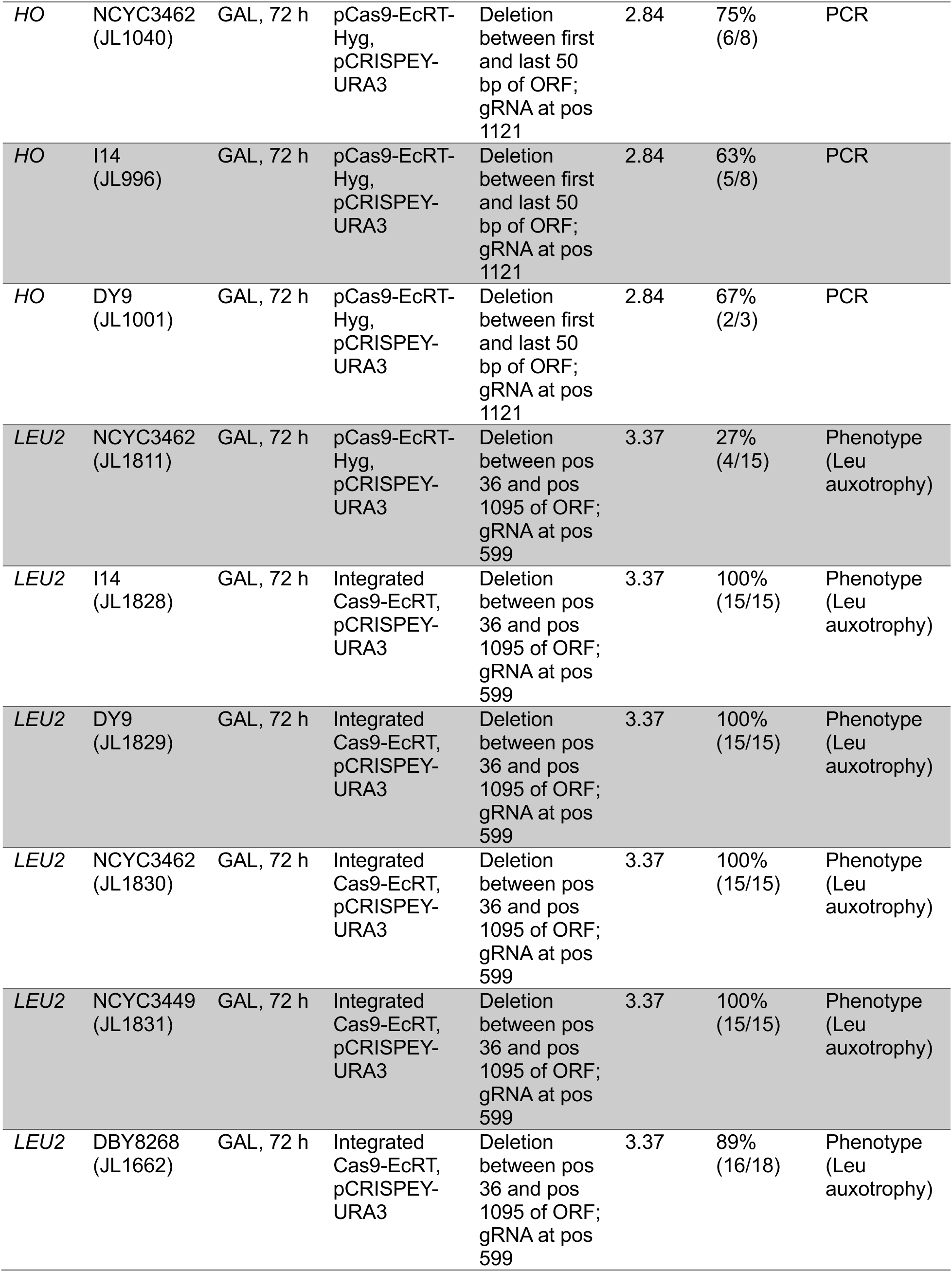
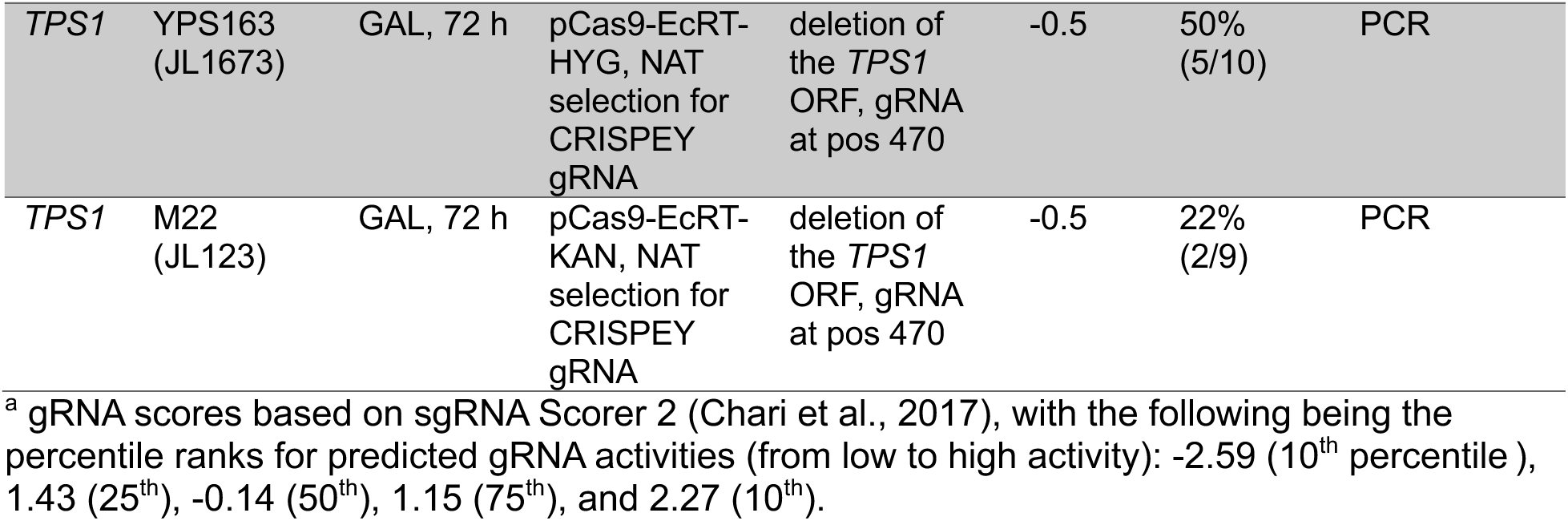
Efficiency of genome editing for other mutations.

We first sought to compare editing efficiencies for genomically integrated versus plasmid-encoded Cas9-EcRT, and also for antibiotic versus auxotrophic selectable markers. We found no significant difference in editing efficiencies for genomically-encoded versus plasmid-encoded Cas9-EcRT, nor for the use of HygMX or NatMX antibiotic resistance markers versus *URA3* or *HIS3* auxotrophic markers for the plasmid system (Figure 2). In each case, we achieved ∼80% editing efficiency after 24 hours of induction and ∼98% after 48 hours (Figure 2). We next performed every possible pair-wise comparison of antibiotic markers for the Cas9-EcRT and CRISPEY plasmids. The choice of selectable markers had a modest effect on editing efficiencies, with >98% editing efficiencies for all combinations tested at 72 hours of induction (Figure 3), but with a range of 64% to 99% at 48 hours (Figure 3). The combination of the KanMX and HygMX markers had the lowest editing efficiencies (64%-82% editing efficiencies at 48 hours of induction; Figure 3), while the other markers averaged 94% (range of 88% – 99%; Figure 3). Thus, our recommendation is to avoid Hyg-Kan combinations unless necessary, and to increase the editing time to 72 hours when the use of those markers is unavoidable.

**Figure 2.**
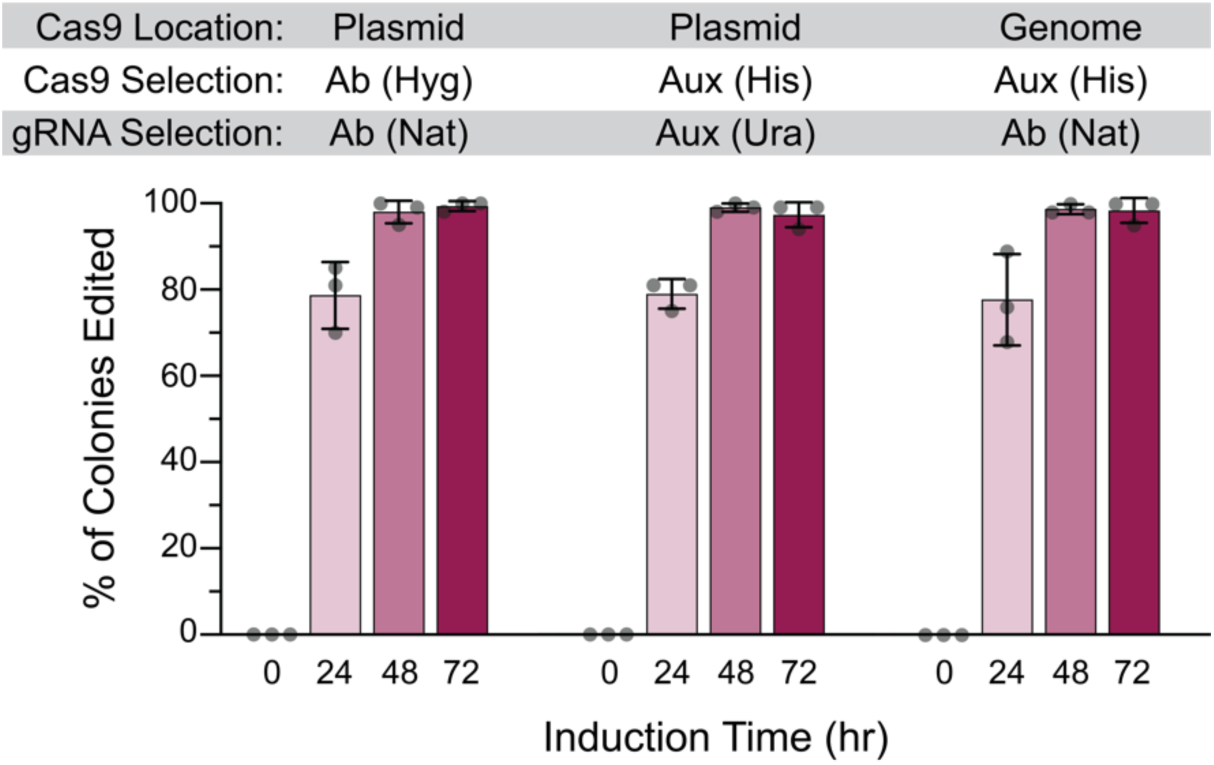
Plasmid-encoded Cas9-RT is as effective for editing as genomically encoded. Cells contained either genomically integrated (at the *his3Δ1* locus) or plasmid-encoded galactose inducible Cas9 and Ec86 retron (pCas9-EcRT-GAL), along with pCRISPEY-GAL-ADE2 containing a gRNA and repair template that introduces a frameshift mutation into *ADE2*. Cells were passaged for the indicated amount of time on galactose-containing selective media and then plated onto YPD to score editing efficiency. Error bars denote the mean and standard deviation of biological triplicates.

**Figure 3.**
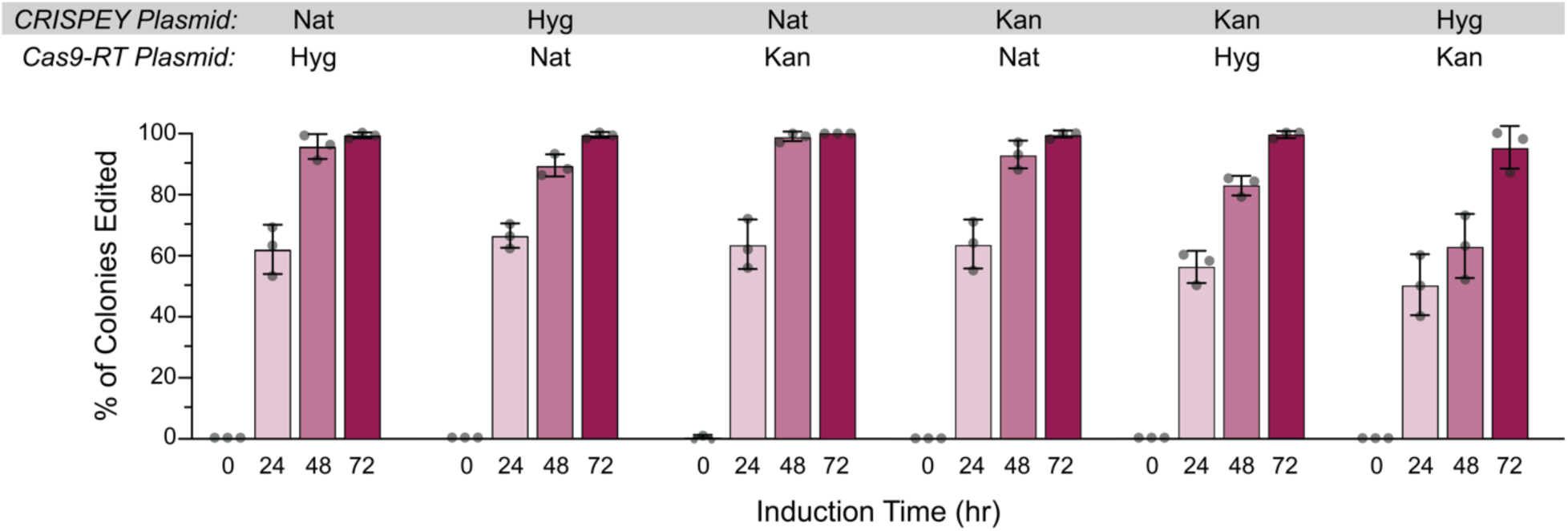
The choice selectable marker has a small effect on editing efficiency. Cells were transformed with galactose-inducible pCas9-EcRT-GAL and pCRISPEY-GAL-ADE2 plasmids containing the denoted antibiotic resistance markers. Cells were passaged for the indicated amount of time on galactose-containing selective media and then plated onto YPD to score editing efficiency. Error bars denote the mean and standard deviation of biological triplicates.

### High editing efficiencies in generating homozygous diploids

Because wild strains are generally diploid, we tested editing efficiencies in a panel of strains that were either diploid or haploid. Notably, *ade2* mutations are recessive, so only homozygous *ade2*/*ade2* mutants have red colonies. For this experiment, we included the reference lab strain (S288C derived) used in all previous experiments, along with a wild oak strain (YPS163), a wild vineyard strain (M22), and a clinical strain (YJM339). There were strain-specific differences in editing efficiency, with the lab strain S288c having the highest editing efficiencies across both diploid and haploid panels, and the clinical isolate having the lowest efficiencies (Figure 4). Even for the clinical strain however, 91% editing efficiencies were observed in diploids after 72 hours of induction, compared to ∼99% editing efficiencies for the each of the other diploids under the same conditions. While we have not tested this, the high editing efficiencies seen in diploids suggest that this method could be used to generate homozygous edits in polyploid commercial strains of yeast (e.g., brewing yeast) that often cannot sporulate (Gallone et al., 2016) and have thus been challenging for genetic analysis.

**Figure 4.**
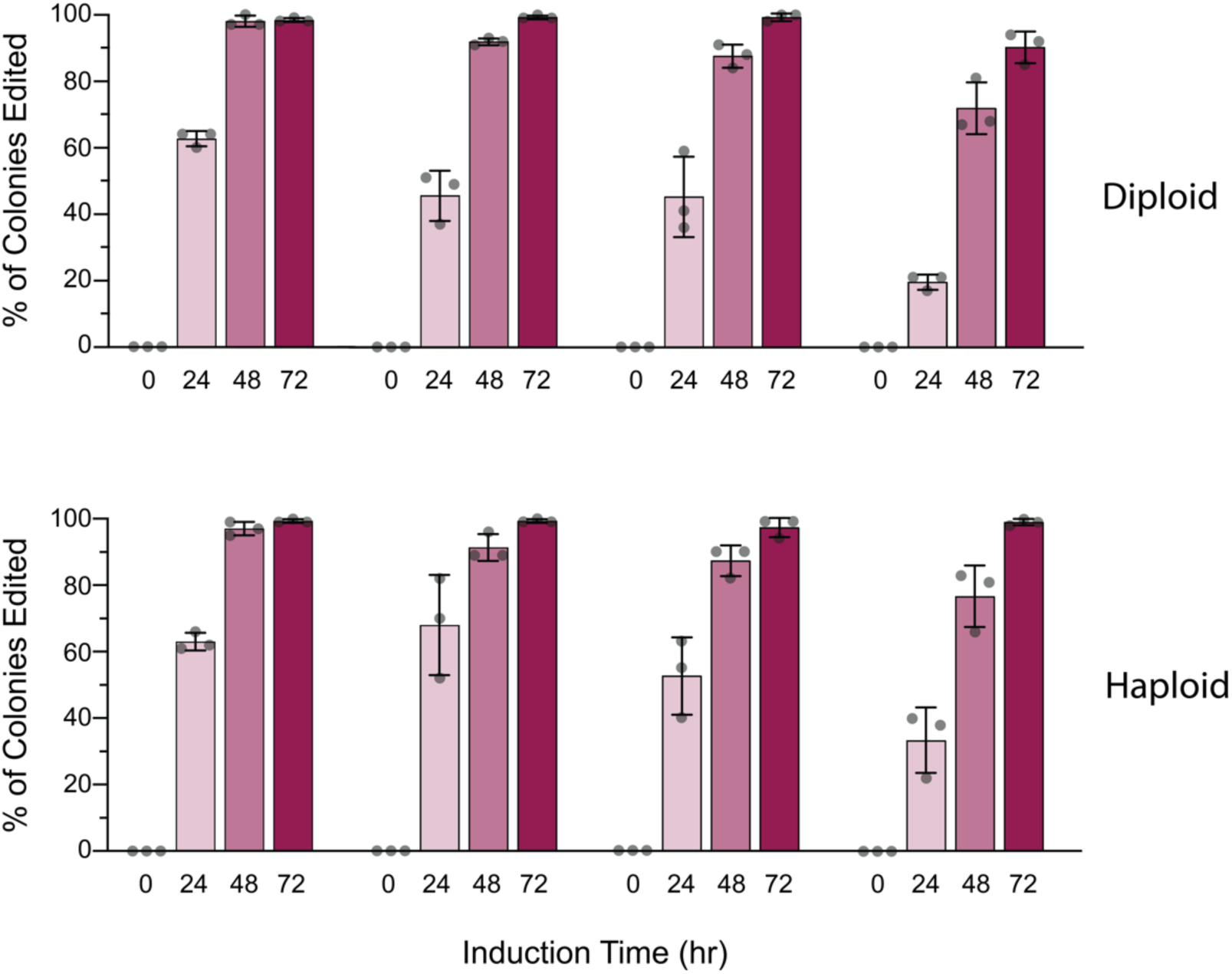
High editing efficiencies in wild diploid yeast. Diploid or haploid derivatives of a lab strain (S288C-derived DBY8268), wild oak (YPS163), vineyard (M22), and clinical (YJM339) were transformed with galactose-inducible pCas9-EcRT-GAL-Hyg and pCRISPEY-GAL-Nat-ADE2 plasmids containing the denoted antibiotic resistance markers. Cells were passaged for the indicated amount of time on galactose-containing selective media and then plated onto YPD to score editing efficiency. Error bars denote the mean and standard deviation of biological triplicates.

### High efficiency genome editing with a β-estradiol-inducible system

One of our main motivations for constructing these plasmids was to use them with fully prototrophic wild yeast strains. However, some strains grow poorly in galactose (Palme et al., 2021), making them incompatible with the original CRISPEY system. Thus, we created a second version of this system by swapping the *GAL* promoter with one inducible by β-estradiol. We chose the Z_3_EV system (McIsaac et al., 2013), which consists of an artificial transcription factor (TF) (with 3 zinc finger domains, hence the Z_3_ name) and a synthetic promoter (pZ_3_EV) with DNA-binding sites for the artificial TF. The artificial TF consists of DNA-binding domains from the mouse Zif268 TF fused to the human estrogen receptor and the VP16 activation domain. In the absence of β-estradiol inducer, the estrogen receptor domain interacts with the Hsp90 chaperone complex, which conceals a nuclear localization signal. The chaperone complex is displaced by β-estradiol binding to the estrogen receptor, thus allowing nuclear localization, binding to cognate DNA-binding sites, and gene activation. A benefit of this system is that β-estradiol can be used as the inducer with any carbon source, allowing for gene induction with glucose. One caveat though, and a reason why galactose induction is still the ‘gold standard’ is that the Z_3_EV system results in lower induction than the galactose inducible system (Arita et al., 2021).

We tested the β-estradiol inducible system in different media: SC galactose, SC dextrose, and YPD. Editing efficiencies were high (98-99% after 48 hours of induction) when performed on SC galactose and YPD, and slightly lower for SC glucose (88%, Figure 5 and Figure S1), strongly suggesting that the potentially lower expression of the Z_3_EV a limiting factor. There was considerable editing that occurred prior to induction (10-23%), suggesting ‘leaky’ expression in the absence of β-estradiol, so the system may not be ideal if the edit confers a significant fitness defect. Nonetheless, editing efficiency with β-estradiol was at least as high as for galactose-induction, and thus constitutes an effective alternative system for strains that grow poorly on galactose.

**Figure 5.**
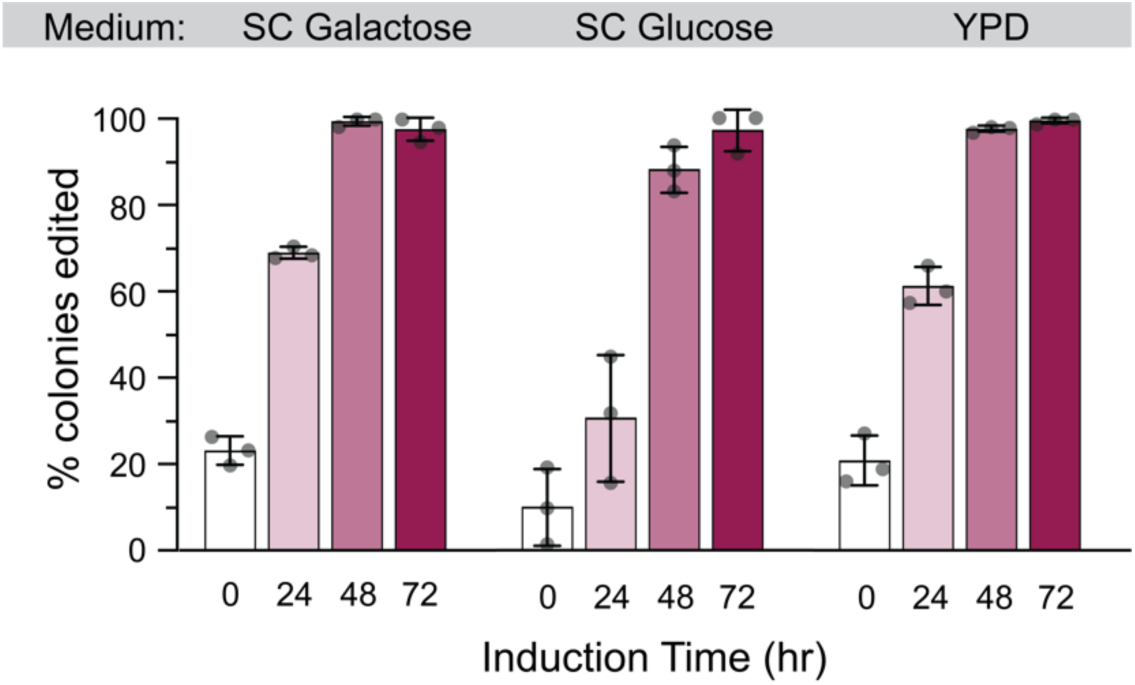
High editing efficiencies with an estradiol-inducible CRISPEY system. Cells were transformed with β-estradiol-inducible pCas9-EcRT-Z3 and pCRISPEY-Z3-ADE2 plasmids containing the denoted antibiotic resistance markers. Cells were passaged for the indicated amount of time on β-estradiol-containing selective media and then plated onto YPD to score editing efficiency. Error bars denote the mean and standard deviation of biological triplicates.

### Efficient generation of diverse deletion and point mutations

Our lab has been using the dual plasmid system to generate deletions and point mutations in diverse strains (Table 3). For deletions, the repair template contains 50-bp each of the 5’ and 3’ regions flanking the region to be deleted. For deletions, our overall efficiency is 62% (range of 34% – 86%), which likely depends greatly on the strain and genomic context (Table 3). This is lower than the editing efficiency for point mutations across all of our experiments (>92% at 48 hours including the clinical strain) and likely reflects the distance between the homologous arms of the repair template. However, because deletions can easily be screened via PCR, the generation of markerless deletions at even the lower end of that range makes the CRISPEY system a valuable part of the yeast deletion toolbox. For point mutations, having high editing efficiencies that approach 100% in many cases dramatically decreases the amount of sequencing needed identify correctly editing strains. The resulting strains are easily cured of plasmid following 24 – 48 hours of passaging on non-selective media.

### A protocol for use of the dual plasmid system

Below, we provide a protocol for groups who wish to implement the method in their own labs. The workflow includes 1) design of an oligonucleotide containing the gRNA and repair template, 2) cloning of the oligonucleotide into a pCRISPEY vector, 3) simultaneous transformation of the pCRISPEY and pCas9-EcRT plasmids, 4) induction of the genome editing, 5) screening of clones for desired edits, and 6) ‘curing’ of the plasmids. Below, we provide details for each step:

*Step 1: design of oligoncucleotide containing the gRNA and repair template*. The oligonucleotide is 190-nt in length and can be ordered from a number of commercial vendors (we use Integrated DNA Technologies (IDT)).The oligonucleotide consists of a 5’ invariant sequence for primer binding, the repair template with at least 50-bp homology on each side flanking the targeted edit, an invariant retron sequence that is necessary for reverse transcription of the repair template into ssDNA, the gRNA sequence, and then a final invariant sequence for 3’ primer binding. Thus, only the repair template and gRNA need to be designed for each new construct. We use CRISpy-pop (Stoneman et al., 2020) for gRNA design, which has an easy web interface (https://crispy-pop.glbrc.org/), and can be used to rank gRNAs for 1,011 published yeast genomes (Peter et al., 2018). CRISpy-pop also has the benefit of using sgRNA Scorer 2.0 (Chari et al., 2017) to predict Cas9 activity for gRNA, as well as Cas-OFFinder (Bae et al., 2014) to identify potential off-target matches. Ideally, gRNAs should have an activity score of at least zero (∼50^th^ percentile for expected activity) and should be as close to possible to the nucleotide(s) to be edited. The repair template should then include the gRNA sequence and should ideally edit the PAM sequence to eliminate cutting by Cas9 following successful homologous recombination. For generation of missense mutations, if editing of the PAM sequence does not achieve the desired allele, our recommendation is to introduce both the desired edit and a silent mutation of the PAM site (using an alternative codon with similar usage to that of the wild-type codon). In that case, a strain containing the silent mutation edit alone should be constructed as a control strain for comparisons to the mutant(s) of interest. For generation of deletions, the repair template simply should have 50-nt each on the 5’ and 3’ regions that flank the desired region to be deleted, with the PAM located somewhere within the region to be deleted.

*Step 2: cloning of the gRNA-repair template oligonucleotide into a pCRISPEY vector.* Cloning efficiencies for introducing the 20-nt gRNA oligonucleotide have generally been low (Antony et al., 2022), but in our hands, Gibson cloning of the gRNA-repair template oligonucleotide typically achieves cloning efficiencies of >90%. The oligonucleotide is amplified with CRISPEY_F and CRISPEY_R primers that add 40-bp of vector sequence for Gibson assembly, using the following cycling conditions:

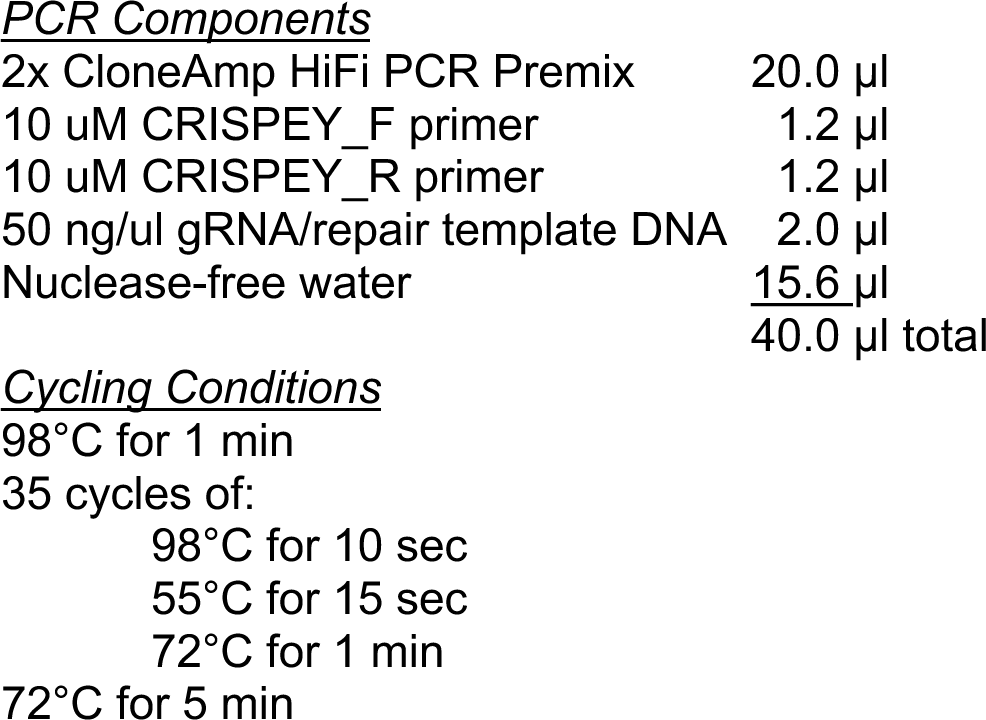

The pCRISPEY plasmid is digested with either *Not*I (GAL inducible) or *Xho*I (β-estradiol inducible) to linearize the vector. Both the linearized vector and the gRNA-repair template product are purified and fused together using Gibson cloning. We use the NEBuilder® HiFi DNA Assembly kit according to the manufacturer’s protocol and then transform 2 µl of each reaction into 50 µl of chemically competent *E. coli* cells. We generally purify plasmid from 4 transformants and performing Sanger sequencing using the CRISPEY Retron SEQ F primer.

*Step 3: simultaneous transformation of the pCRISPEY and pCas9-EcRT plasmids into yeast.* Yeast are transformed with both the pCRISPEY and pCas9-EcRT plasmids via the high-efficiency yeast transformation method of (Gietz & Schiestl, 2007). For wild yeast strains and antibiotic selection, higher transformation efficiencies are obtained by first plating on YPD and letting cells outgrow for 24 hours before replica printing to antibiotic plates.

*Step 4: induction of the genome editing.* For the GAL-inducible system, transformants are inoculated into selective media (antibiotic or auxotrophic) containing 2% raffinose as a non-GAL repressing carbon source and grown 24 hours at 30°C with shaking. Sixty µl of the culture is inoculated into 1.94 ml selective induction media containing 2% galactose (SC-GAL or YP-Gal) to begin induction. Following 24 hours of induction, 60 µl of culture is sub-cultured into 1.94 ml of induction media for a total of 48 hours of induction. A third sub-culture for a total of 72 hours of induction can be used for particularly recalcitrant edits. Seventy-two hours of induction is recommended if uses the Nat/Hyg combination of markers.

For the β-estradiol inducible system, cells can be grown in selective media containing dextrose (or any carbon source), and induction is started by adding 60 µl of cells to 1.94 ml of selective induction media (SC or YPD) containing 1 µM β-estradiol (β-estradiol can be stored at −20°C as a 5 mM stock in 50% ethanol).As for the GAL-inducible system, 60 µl of culture is sub-cultured into 1.94 ml of induction media following 24 hours of induction for at total 48 hours of induction.

*Step 5: screening of clones for desired edits.* Following 48 or 72 hours of induction, cells are streaked to selective media (e.g., YPD plus antibiotics) and incubated for 2-3 days at 30°C until colonies are visible. Isolated colonies can be screened for successful editing by PCR (in the case of deletions) or sequencing. We generally sequence 4 isolates for missense mutations because editing efficiencies tend to be >90%. The efficiency for deletions tends to be lower, so we screen 10-20 via diagnostic PCR. For challenging edits, efficiencies may be higher if plating cells onto selective instead of non-selective media (likely due to continued editing occurring on plates, Figure S2), though this may also increase the amount of time necessary to ‘cure’ the plasmids during the next step.

*Step 6: ‘curing’ of the plasmids.* Strains with the correct genotype are inoculated into non-selective media (e.g., YPD) and grown for 24-48 hours. Cells are then streaked onto non-selective plates, and ∼20 colonies are patched onto plates containing each antibiotic to identify sensitive isolates that have lost both plasmids. We generally obtain multiple colonies that have lost both plasmids at this point, but one can sequentially cure each plasmid if the rate of loss is too low.

### Caveats and Considerations

The CRISPEY system relies on homologous recombination, thus, mutants deficient in homologous recombination cannot be directly edited. If one wishes to generate targeted edits in a strain deficient in homologous recombination, one could generate the edits in a wild-type strain and then mate with the strain with the desired background and sporulate to isolate the genotype of interest. Based on our experiments, we found that editing was lower using vectors with Nat and Hyg resistance in combination. In general, the editing efficiencies are still quite high (>95%) after 72 hours of induction with that combination, though we would recommend using a combination of Nat, Kan, and/or *URA3* markers, especially for gRNAs predicted to have lower efficiency. Multiplex editing should be possible by combining the use of pCRISPEY plasmids with different markers, though we did not test that here. Our recommendation for multiplexing would be to increase the number of rounds to induction (72 hours or 96 hours) to increase the frequency of obtaining clones with all of the desired edits. However, because the CRISPEY system is markerless, any number of desired mutations can be generated sequentially.

### Conclusions

We have created a panel of CEN plasmids that allow for efficient retron-mediated genome editing in fully prototrophic wild yeast strains with a minimal number of steps. The system requires ordering of only a single 190-nt oligonucleotide consisting of the gRNA and repair template to be cloned into the pCRISPEY vector. The Cas9-EcRT and pCRISPEY plasmids can be simultaneously transformed into a yeast strain of interest, and generally 48 hours of induction is sufficient for >90% editing efficiencies for point mutations across a diverse panel of strain backgrounds. The Cas9-EcRT and pCRISPEY plasmids can be cured by passaging over 24 – 48 hours, thus completing the markerless editing. The system can also be used to generate homozygous edits in diploids with high efficiency (>97% following 72 hours induction), and we also introduce a by β-estradiol inducible system as an alternative for strains that grow poorly on galactose. Overall, this system simplifies high-efficiency targeted genome editing in yeast.

## Acknowledgements

This work was supported in part by National Science Foundation Grant MCB-1941824 (JAL) and the Intramural Research Program of the National Human Genome Research Institute, NIH (1ZIAHG200401) (MJS). JW was supported by the Arkansas IDeA Network of Biomedical Research Excellence (INBRE) Summer Research Fellowship program, which is funded through NIH-NIGMS grant P20 GM103429. The funders had no role in study design, data collection and interpretation, or the decision to submit the work for publication.

---

**Supplemental Figure 1.**
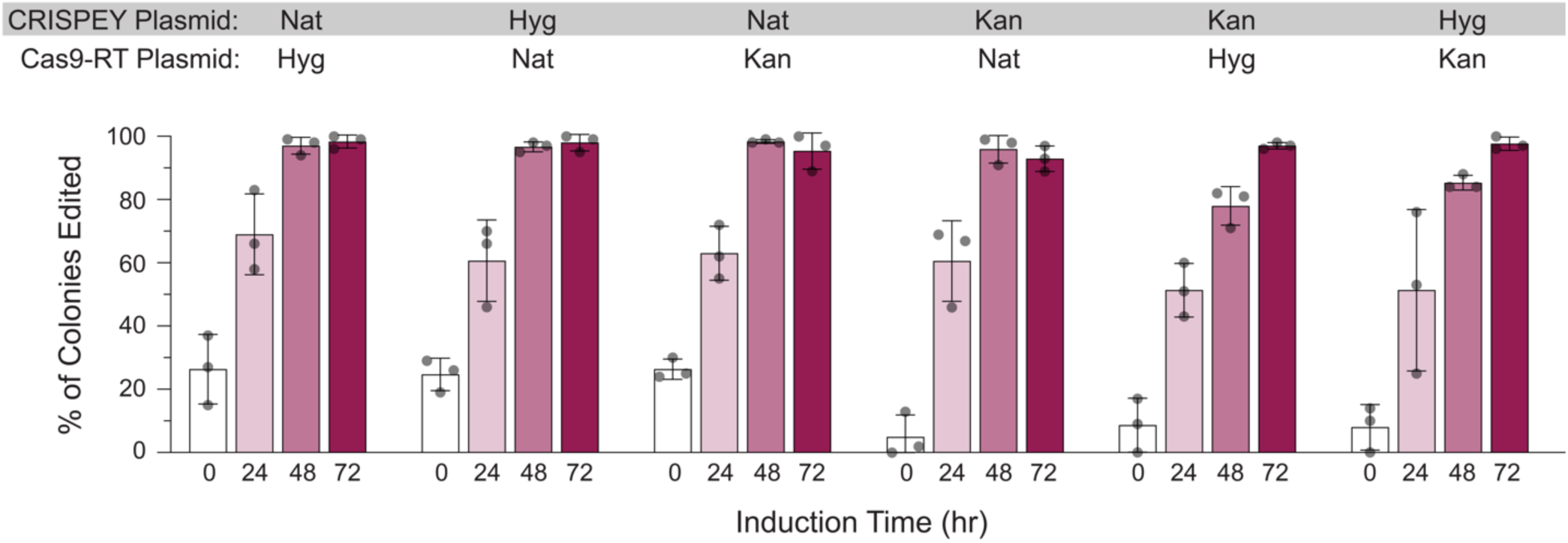
The choice selectable marker has a small effect on editing efficiency with the estradiol-inducible CRISPEY system. Cells were transformed with β-estradiol-inducible pCas9-EcRT-Z3 and pCRISPEY-Z3-ADE2 plasmids containing the denoted antibiotic resistance markers. Cells were passaged for the indicated amount of time on β-estradiol-containing selective media and then plated onto YPD to score editing efficiency. Error bars denote the mean and standard deviation of biological triplicates.

**Supplemental Figure 2.**
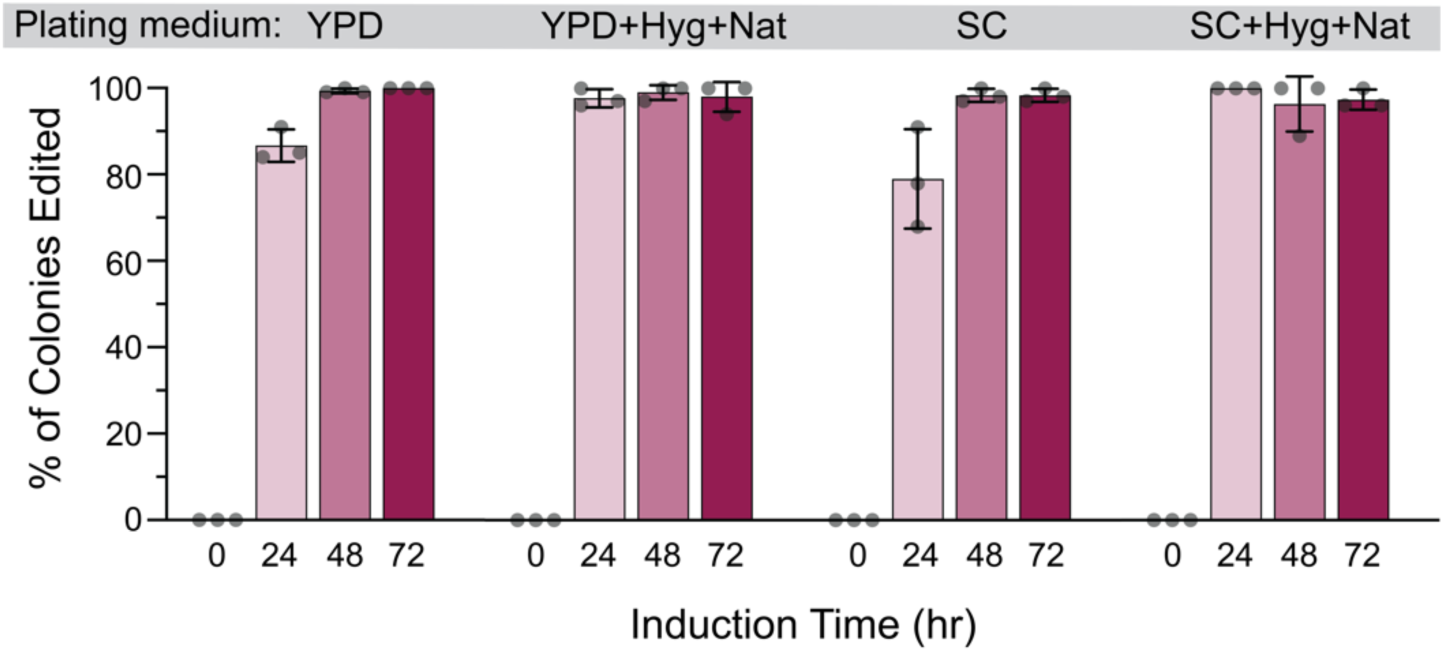
Plating medium has little effect on editing efficiency. Cells were transformed with galactose-inducible pCas9-EcRT-GAL and pCRISPEY-GAL-ADE2 plasmids containing the denoted antibiotic resistance markers. Cells were passaged for the indicated amount of time on galactose-containing selective media and then plated onto either non-selective (YPD or SC) media or selective media (YPD or SC plus antibiotics) as indicated to score editing efficiency. Error bars denote the mean and standard deviation of biological triplicates.

**Table S1:**
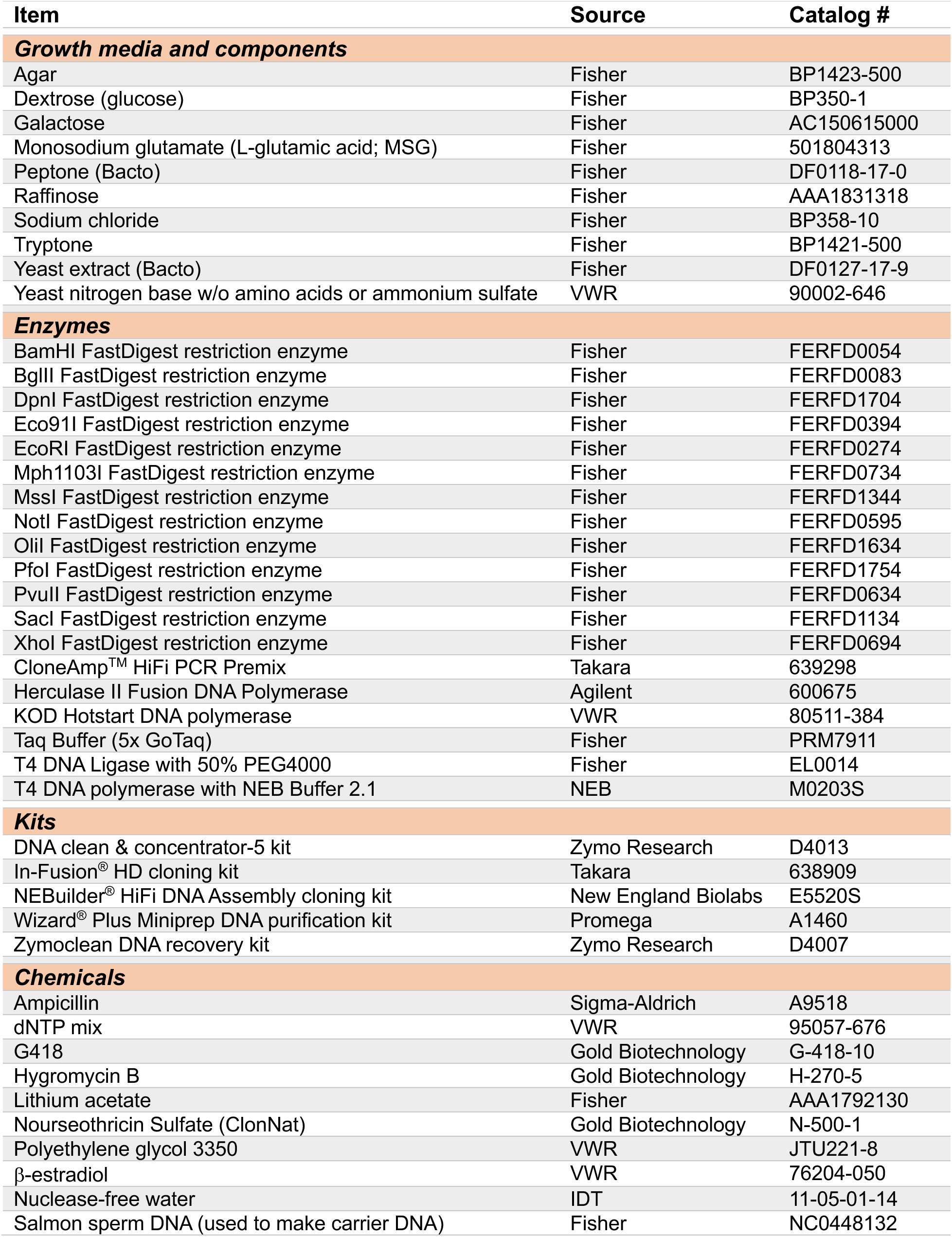
Reagent, media and chemicals used in this study.

**Table S2:**
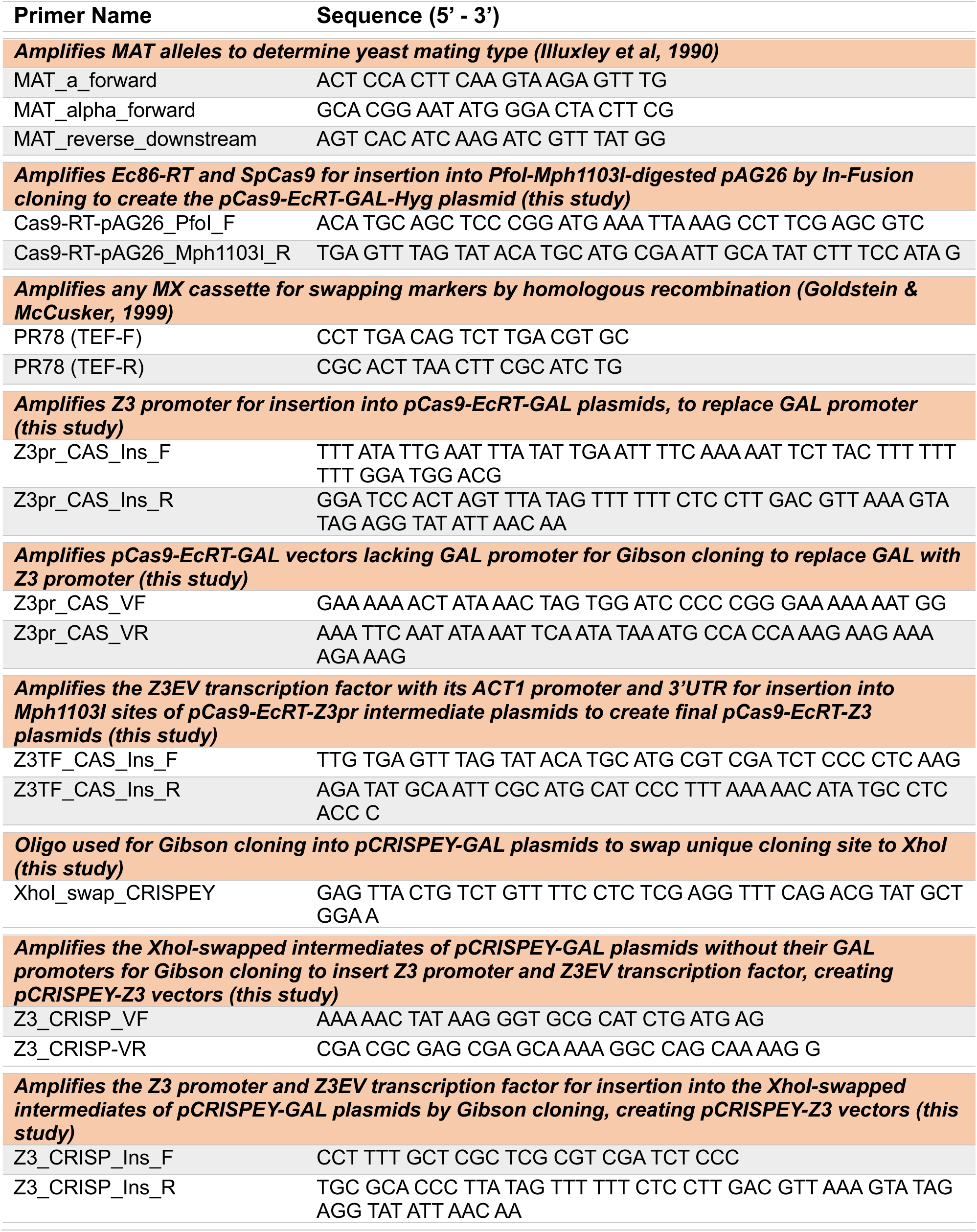
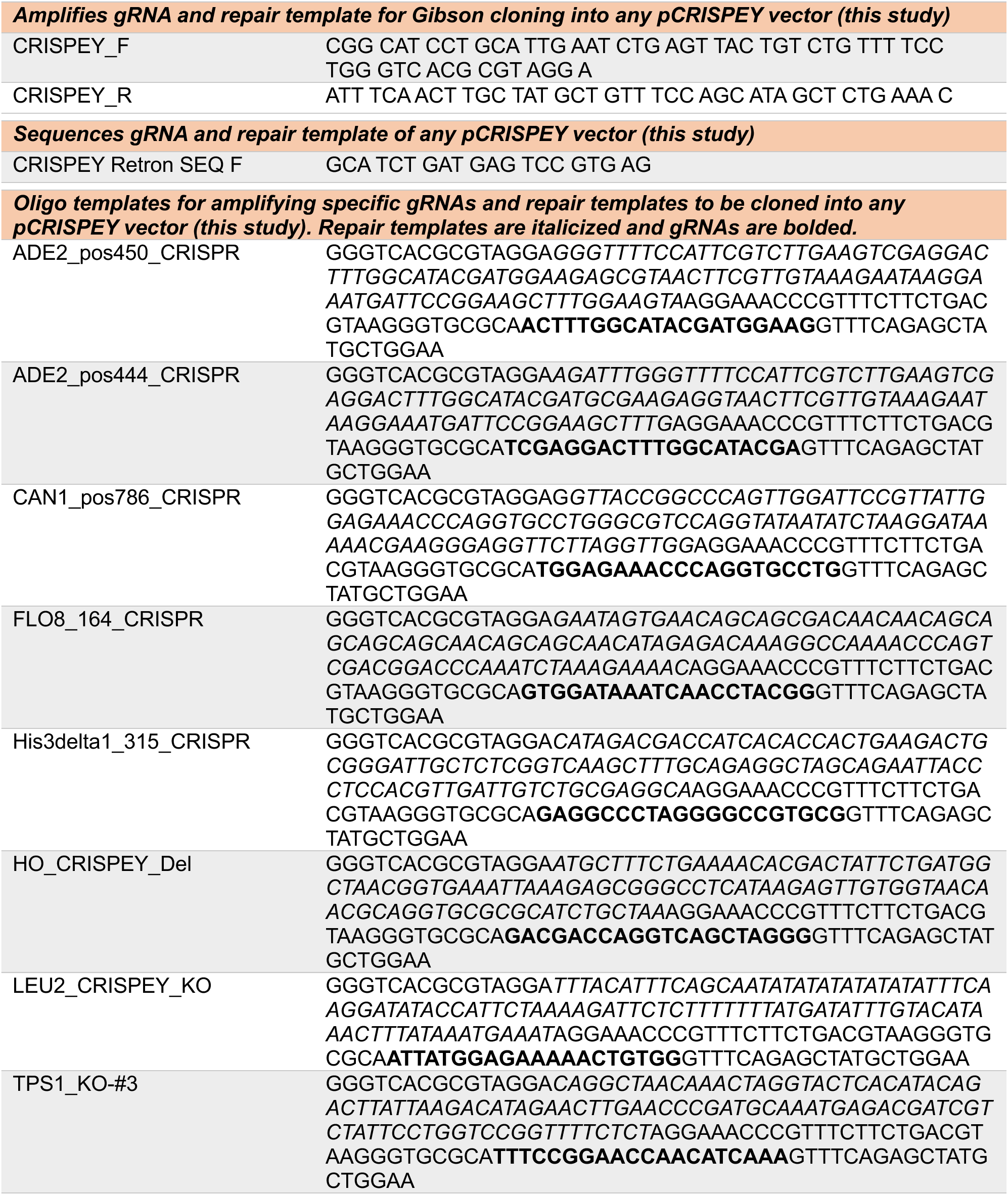
Primers used in this study.

**Table S3:**
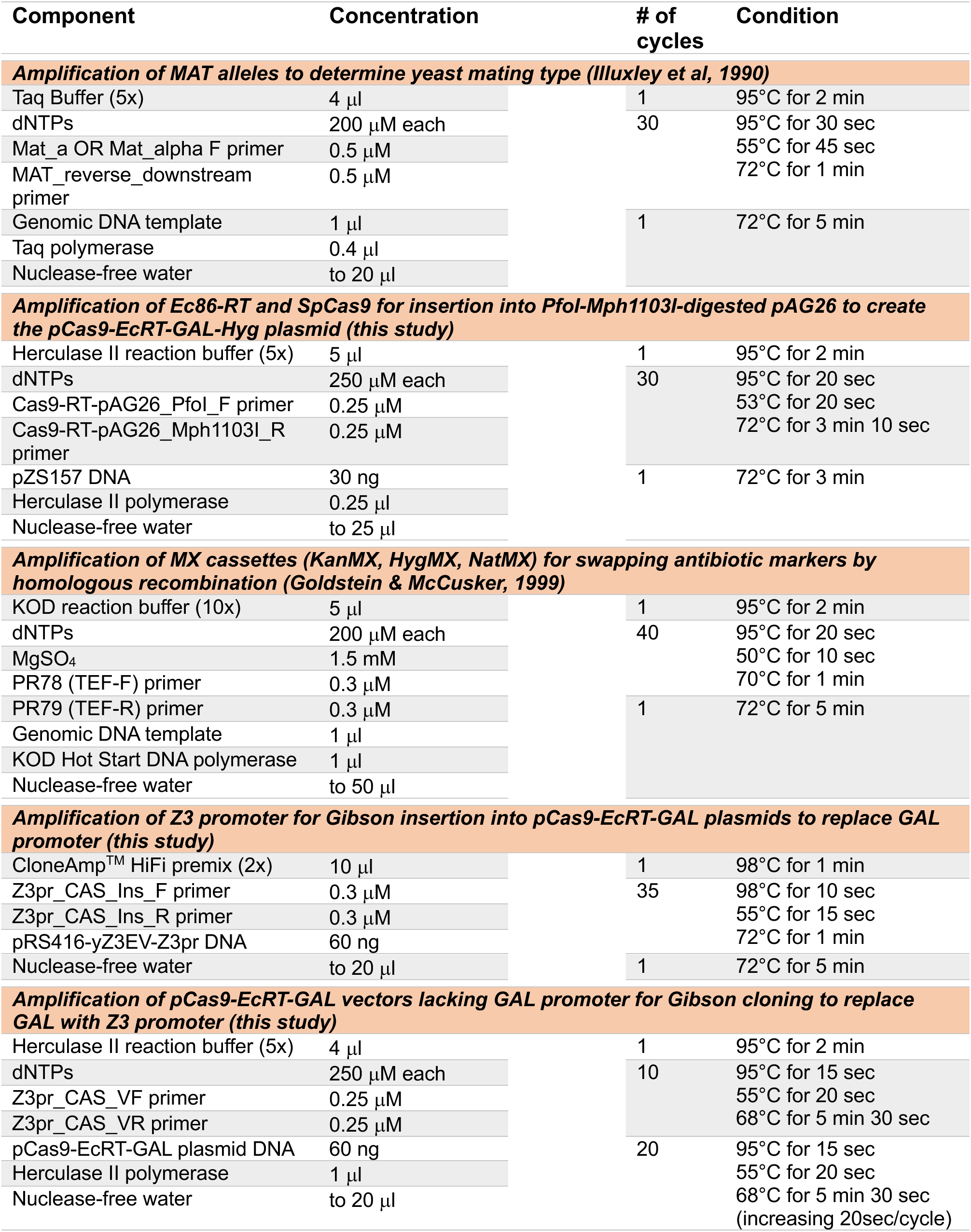
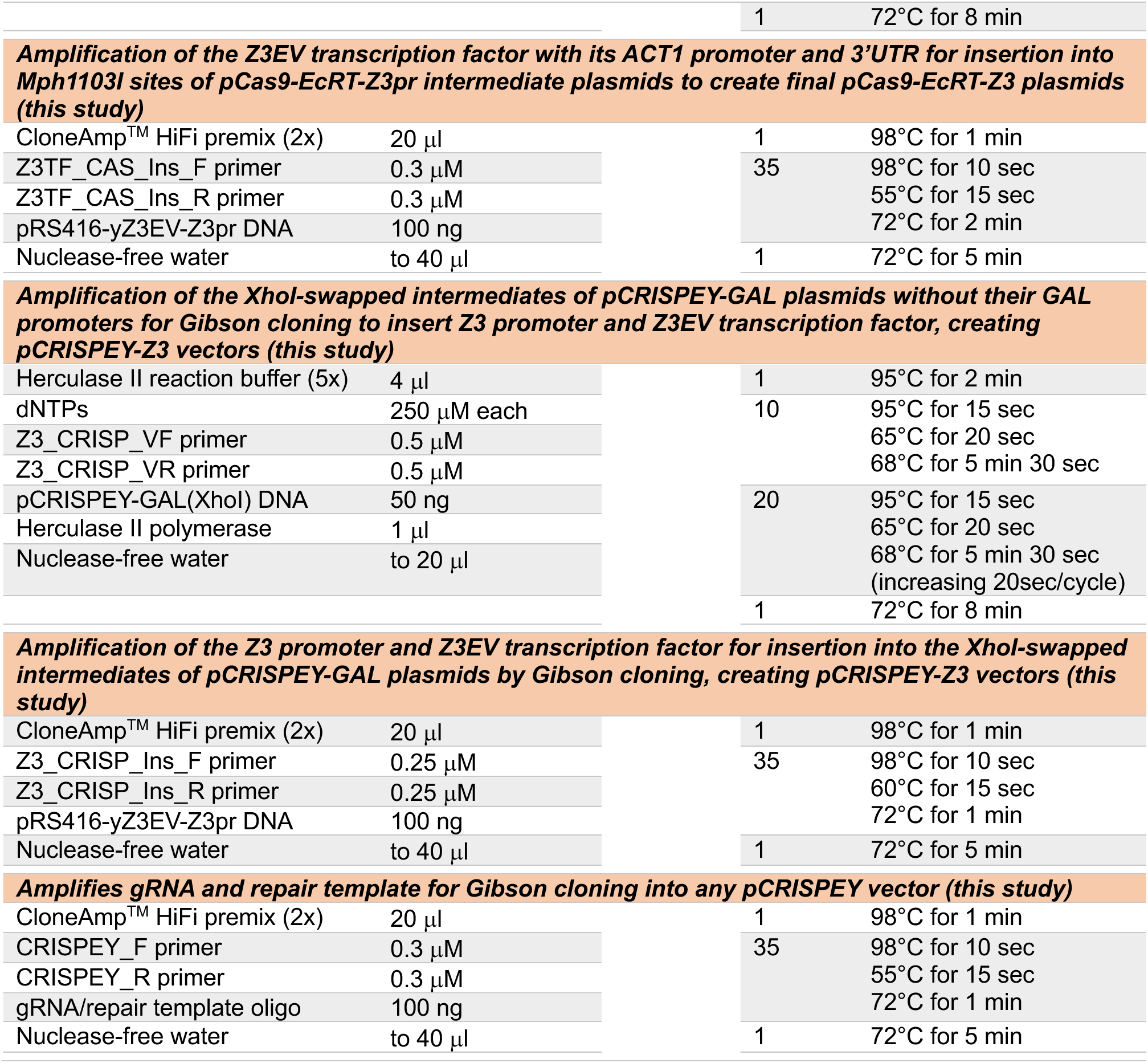
PCR conditions for all amplifications from this study.

## Supplemental Files

**File S1: Sequences of pCRISPEY and pCas9-EcRT plasmids.**

